# CellTouch: Learning to separate touching nuclei in anisotropic 3D cell cultures

**DOI:** 10.64898/2026.11.27.769674

**Authors:** Istvan Grexa, Marton Bolf, Emese Recsanyi, Csaba Molnar, Akos Diosdi, Peter Horvath

**Affiliations:** Synthetic and Systems Biology Unit, HUN-REN Biological Research Centre, Szeged, Temesvári körút 62, Szeged 6726, Hungary; Single-Cell Technologies Ltd, Temesvári körút 62, Szeged 6726, Hungary; Doctoral School of Computer Science, University of Szeged, Árpád tér 2, Szeged 6720, Hungary; Doctoral School of Biology, University of Szeged, Közép fasor 52, Szeged 6726, Hungary; Institute of AI for Health, Helmholtz Zentrum München, Ingolstädter Landstraße 1, Neuherberg 85764, Germany; Institute for Molecular Medicine Finland (FIMM), University of Helsinki, Tukholmankatu 8, Helsinki 00290, Finland

**Keywords:** Anisotropic segmentation, 3D cell cultures, 3D segmentation, AI

## Abstract

3D-oids, such as spheroids and organoids, are essential models in drug discovery and precision medicine, and quantitative analysis requires every nucleus to be segmented separately. Accurate 3D instance segmentation remains challenging, because nuclei are densely packed and touching, and volumes are sampled more coarsely along the Z-axis than laterally. Most existing methods were developed for near-isotropic data and tend to merge neighbouring nuclei. To address this challenge, we propose CellTouch, a deep learning method for anisotropic 3D nuclei instance segmentation that predicts flow fields and a contact channel, which separates touching nuclei directly from the network output. Ablations showed that the contact channel is the key design element, and CellTouch outperformed state-of-the-art methods on in-house data and widely used public datasets. Most importantly, dense Z-sampling is not required: CellTouch segmented nuclei as accurately at a 2 µm Z-step as at 0.5 µm, reducing acquisition time and data volume four-fold.

## Introduction

Recent advancements in high-content screening (HCS) and single-cell analysis showed that morphology is now a key readout, complementing molecular profiling techniques like omics (e.g. genomics, proteomics, and transcriptomics) and spatial information [1]. Morphology not only helps to describe phenotypes, but extracted morphological features can also capture the cellular states, environmental interactions, and perturbation responses [2]. As a result, rapid adoption of large-scale imaging assays and morphological reference datasets has been established for two-dimensional (2D) cell cultures and tissue sections where single-cell segmentation and characterisation underpin the analysis [3]. However, the generation of large-scale imaging datasets alone is insufficient without appropriate analytical tools. Consequently, initiatives such as cell atlases and morphology profiling consortia have accelerated both the creation of imaging resources and the development of computational methods for 2D segmentation [4], [5].

3D-oids (e.g., spheroids, organoids, assembloids, and connectoids) are promising model systems emerging as an alternative because these models better recapitulate tissue architecture, cellular heterogeneity, and cell-cell interactions than conventional 2D cultures, making them increasingly valuable for toxicology, regenerative medicine, and translational research [6], [7], [8]. In order to introduce and integrate these models into preclinical pipelines, proper characterisation, validation, and qualification are necessary [9]. Needless to say, to fully utilise large-size and complex 3D-oid models, the combination of microscopy image-based validation and high-accuracy segmentation methods is essential.

To validate 3D-oids, high-resolution imaging conditions are necessary for visualising their inner parts and characterising not only the model morphology but also their cellular composition, spatial organisation, and phenotypic diversity at single-cell resolution [6]. As a standard, most of the pipelines are built around high-throughput confocal systems, however, as an alternative, light-sheet fluorescence microscopy (LSFM) offers single-cell resolution with fast image acquisition, high penetration depth, and, more recently, the benefits of screening large numbers of samples [6], [10]. Regardless of whether data are acquired using confocal or light-sheet microscopy, current imaging technologies generally produce volumetric datasets with substantially lower resolution along the Z-axis than within the x-y plane. This inherent anisotropy necessitates the development of segmentation and analysis frameworks that can effectively account for unequal spatial sampling across dimensions. Addressing this challenge is essential for the accurate characterisation of 3D biological structures and for the extraction of realistic morphological features from microscopy data.

Thus, many research fields are moving toward fundamentally 3D analyses to routinely generate volumetric datasets at cellular resolution [11], [12]. Analysing such intricate 3D features with current state-of-the-art 2D methods falls short. Capturing the complex 3D structure of biological systems requires advanced technological developments, which leads us to 3D segmentation [13], [14].

Deep learning methods have become the dominant paradigm for 3D microscopy image segmentation, enabling automated analysis of increasingly complex volumetric datasets. Early architectures such as 3D U-Net [15] established the foundation for volumetric semantic segmentation by learning hierarchical image features directly from 3D image volumes. However, semantic segmentation alone is insufficient for many biological applications, where individual cells or nuclei must be separated as distinct instances. To address this challenge, several specialised instance-segmentation methods have been developed. Cellpose [16] introduced a flow-field representation in which pixels are assigned vectors pointing towards object centres, enabling robust segmentation across diverse cell types. However, it lacks a true 3D mode since 3D segmentation is obtained by combining 2D predictions from orthogonal slices or by stitching 2D masks across adjacent slices, and elongated objects tend to be over-segmented. Omnipose [17] extends the Cellpose paradigm by directing flows towards the object’s medial axis rather than its centroid, improving segmentation of elongated and irregularly shaped cells. However, a well-known limitation of the method is separating touching objects. StarDist-3D [18] instead represents objects as star-convex polyhedra, achieving strong performance on nuclei with approximately star-convex shapes. However, this geometric constraint may limit the accurate representation of irregular nuclear morphologies, while the finite number of rays further restricts the fidelity of reconstructed object boundaries [19]. Recent benchmarks have also highlighted substantial performance variability across datasets [14]. More recently, NucVerse3D [20] combined a residual attention 3D U-Net with centroid-directed gradient fields to perform volumetric instance segmentation of nuclei.

Despite these advances, accurate segmentation of densely packed biological samples remains challenging. Many methods assume relatively regular object geometries or require volume resampling to compensate for anisotropic voxel spacing. Furthermore, touching or overlapping nuclei frequently result in merged predictions that must be separated through post-processing strategies such as watershed-based splitting or centroid clustering. Consequently, robust instance segmentation of densely populated and anisotropic 3D microscopy datasets remains an active area of research.

We propose CellTouch, a deep learning method that handles anisotropic data and is directly optimised to separate densely packed, touching nuclei. Besides the foreground and flow fields, the network predicts an engineered contact channel that marks only the surfaces where two nuclei touch. Touching nuclei can thus be separated directly from the network output. In an ablation study, we train four target representations with and without the contact channel, and we benchmark CellTouch against four state-of-the-art methods on three in-house and five external datasets. In addition, we image the same spheroids at several axial resolutions to measure how segmentation accuracy depends on axial sampling. We show that moderate axial sampling gives the same or better accuracy than dense sampling, therefore, fewer optical sections are sufficient, which reduces acquisition time, data storage and annotation effort.

## Materials and methods

### Datasets

#### Cell cultures

The T-47D human breast cancer cell line was obtained from the American Type Culture Collection (ATCC) and cultured in RPMI-1640 (Lonza) supplemented by 10% FBS (EuroClone), 0.2 unit/ml insulin (Gibco, Thermo Fisher Scientific). All media contained 1% Penicillin–Streptomycin–Amphotericin B mixture (Lonza). HeLa Kyoto EGFP-alpha-tubulin/H2B-mCherry cervical cancer cell line was obtained from the Cell Lines Service (CLS-300670, Eppelheim, Germany). Cells were maintained in a HeLa Kyoto medium consisting of DMEM (Lonza, Basel, Switzerland), 10% Fetal Bovine Serum (FBS, Euroclone, Milan, Italy), 2 mM L-glutamine (Lonza), 0.5 mg/ml G418 (Gibco, Montana, United States), and 0.5 µg/ml puromycin (Sigma-Aldrich Chemie GmbH, Steinheim, Germany). Cell cultures were maintained at 37 °C and 5% CO_2_ in a humidified incubator.

#### Spheroid generation and fixation

For spheroid generation, the Cell Floater Dish (SPL Life Sciences Co., Ltd., Pocheon, South Korea), a hydrophobic surface-treated petri dish was used. For each petri dish, 400,000 cells were pipetted with 5 ml cell culture media and incubated for 24 hours at 37 °C in a humidified incubator with 5% CO₂. After 24 hours, cells formed small, spherical aggregates suitable for the experiments. Spheroids were then removed from the petri dish and washed three times using DPBS. After washing, 4% PFA was used for 30 min to fix the samples, then washed again with DPBS 3 times and stored at 4 °C in DPBS until imaging. Before imaging, spheroids were incubated in 0.1% Triton X-100 and stained with 1:200 Flash Phalloidin NIR 647 for 60 min. T-47D spheroids were stained with 1 µg/ml DAPI overnight at room temperature. Before imaging, all spheroids were washed with DPBS three times.

#### Microscopes

Spheroids were screened according to previously published methods [21], [22]. Shortly, spheroids were embedded into low-melting agarose and placed under the Leica TCS SP8 Digital LightSheet (DLS) microscope. For imaging, a 25x/0.95 detection objective with a mounted 2.5 mm mirror device was used to capture fluorescence DLS images, illuminated by the 5x/0.15 objective. Images with a resolution of 2048 × 2048 pixels and a pixel size of 0.144 µm were captured with the sCMOS DFC9000 Leica camera. The same spheroids were imaged multiple times using 0.5, 1, 2, 4, 6 µm distance between the images. As a mounting medium, dH₂O was used for every spheroid. For screening, laser intensity was manually adjusted for each channel (i. e. 405, 488, 552, and 638 nm wavelength) using 200 ms exposure time for each laser (maximum laser intensity 350 mW).

#### In-House datasets

The first dataset (Mono– and Co-culture Spheroids – MaCS) consisted of eleven light-sheet volumes of various spheroid compositions, acquired on a Leica SP8 Digital LightSheet microscope with a 25×/0.95 water-immersion objective. Among the eleven spheroids: six volumes were HeLa Kyoto and MRC-5 co-culture spheroids published by Hollandi et al. [23], [24]; two samples were optically cleared, including T-47D and 5-8F monoculture spheroids published by Diosdi et al. [21], [25]; and the last three unpublished spheroids were small T-47D monoculture. The spheroids ranged from about 110 to 260 µm in diameter and were imaged with a lateral pixel size of 0.14 or 0.29 µm and Z-steps of 1 or 3.7 µm. Three co-culture volumes containing 994 nuclei were held out for testing, and the remaining eight volumes containing 4,671 nuclei were mixed into three training and validation folds of six and two volumes.

The second dataset (Axial Sampling Dataset – ASD) included T-47D (n=2, 74 nuclei) and HeLa Kyoto (n=3, 93 nuclei) monoculture spheroid models where cells form a spherical cluster, providing a clearly defined environment [26]. Five spheroids were each imaged as an anisotropy series, re-acquired at Z-steps of 0.5, 1, 2, 4 and 6 µm, and corresponding annotations were generated for each series. The small size range (< 100 µm) was intentionally selected because these spheroids showed exceptional image quality, enabling manual annotation. This experimental design allowed us to assess model performance across anisotropic imaging conditions with different levels of nucleus recurrence, from images where a nucleus was represented by up to 30 ± 6 consecutive masks (0.5 µm series) to images where the same nucleus appeared in only 2.7 ± 0.7 masks (6 µm series).

The third dataset (Large Spheroid Subregions – LSS) consisted of a single T-47D spheroid (diameter around 300 µm, ∼ 2600 cells) from which two regions of interest were selected for detailed analysis. Due to lateral and axial light scattering inherent to the light-sheet imaging system, two crops were chosen to represent distinct imaging conditions. The first crop captured lateral variations in image quality and contained 186 nuclei, whereas the second extended through the entire spheroid and contained 694 nuclei. Together, these regions encompass a broad range of image qualities and include densely packed, frequently touching nuclei in both the lateral and axial directions, providing a challenging dataset for segmentation and analysis.

#### External datasets

The next three datasets (NucMM [27], BlastoSPIM [28], [29], and C. elegans [28]) are coming from public repositories with independent annotations and different specimens. Such datasets were included to test generalisation across organisms, modalities, and voxel sizes. NucMM-Z [27] provides a 0.1 mm³ electron microscopy volume of nearly a whole zebrafish brain (Fig. 1g), with about 170,000 nuclei. The NucMM-M includes a 0.25 mm³ micro-CT volume of part of a mouse visual cortex, with about 7,000 nuclei (Fig. 1f). BlastoSPIM [29] contains manually curated light-sheet images of mouse embryos. BlastoSPIM 1.0 has 573 images from 31 embryos with 11,708 nuclei (Fig. 1d), and BlastoSPIM 2.0 has 80 late-stage images from 25 embryos with 6,628 nuclei (Fig. 1e). We used the low-SNR test series of both, 60 images with 739 nuclei from BlastoSPIM 1.0 and 3 images with 280 nuclei from BlastoSPIM 2.0. The C. elegans [28] dataset contains 28 confocal volumes of L1 larvae (Leica, 63× oil objective) with 15,122 annotated nuclei (Fig. 1.h). The average volume is 1050 × 140 × 140 voxels, with a voxel size of 0.116 × 0.116 × 0.122 µm. Each external dataset was split into training, validation, and test volumes, and all datasets were combined into three folds for the general model. Fig. 1 shows a representative test volume from each set. In total we used 54 NucMM-Z and 8 NucMM-M subvolumes with 13,830 and 1,205 nuclei and 115 BlastoSPIM 1.0 and 75 BlastoSPIM 2.0 images with 2,016 and 6,246 nuclei.

**Fig. 1:**
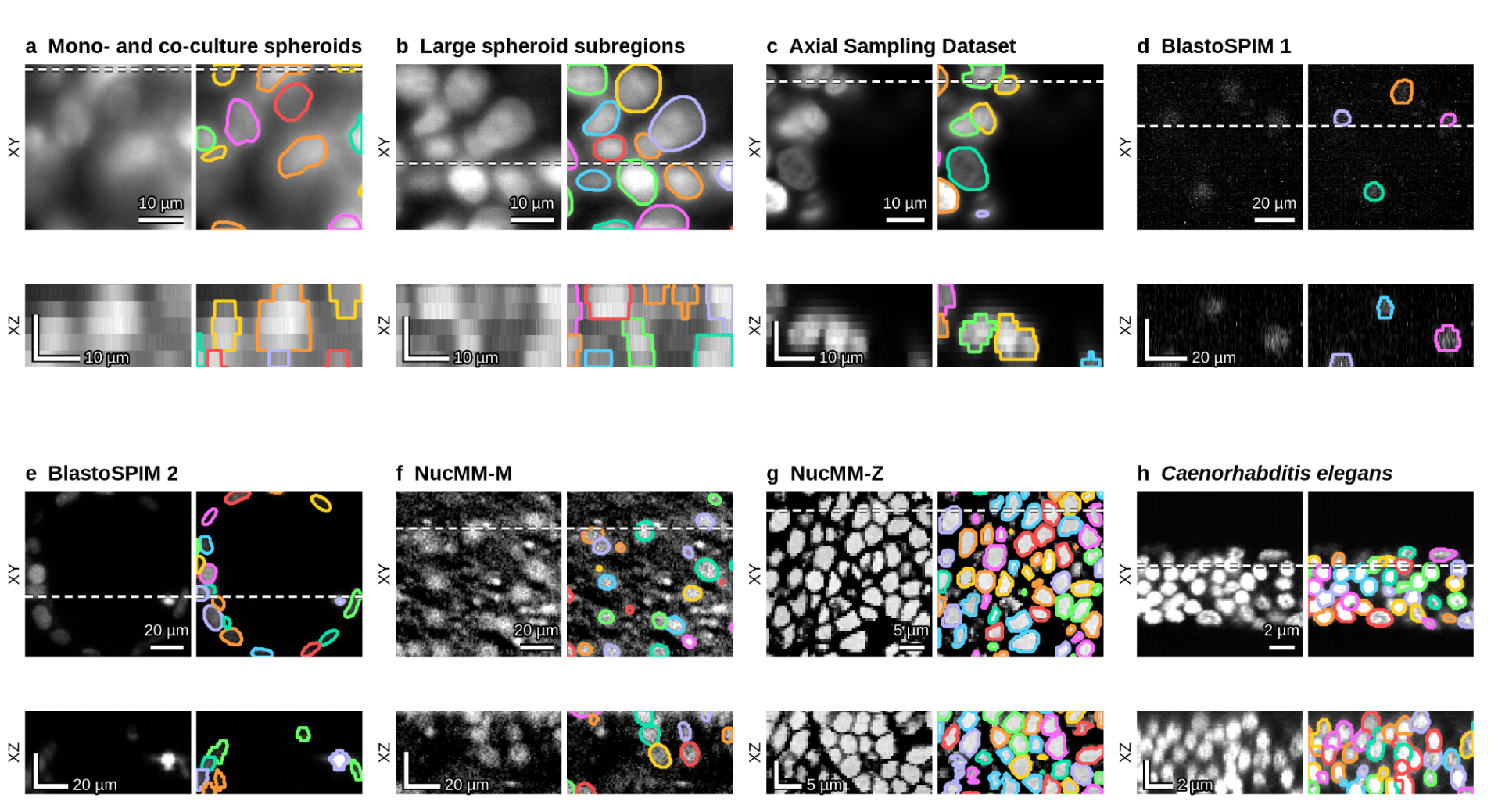
Representative images of the datasets. **a–c**, In-house spheroid test sets: mono– and co-culture spheroids (MaCS; a), large spheroid subregions (LSS; b) and the axial sampling dataset (ASD; c). d–h, External benchmark datasets: BlastoSPIM 1 embryos (d), BlastoSPIM 2 late-stage embryos (e), NucMM-M (f), NucMM-Z (g) and *Caenorhabditis elegans* (h). For each set, the raw image (left) and the ground-truth nuclei outlined, one colour per nucleus (right), are shown in XY (top) and XZ (bottom) views. The XZ view is taken along the dashed line and is drawn at the true physical aspect ratio.

#### Dataset splitting strategy

We trained CellTouch in two regimes. The in-house models were trained only on our own light sheet acquisitions to measure performance within a single imaging setup. The general model was trained on the in-house data together with five public 3D nucleus datasets, *Caenorhabditis elegans*, BlastoSPIM 1.0 and 2.0, NucMM-M and NucMM-Z, to measure how well one model transfers across organisms, microscopes and voxel sizes. Both regimes use the same in-house folds and test sets, so their results on the in-house data can be compared directly.

For training the in-house models, we used a set of spheroid co-cultures and monocultures. Three co-culture volumes, containing 994 nuclei, were held out for testing. The remaining eight volumes, containing 4,671 nuclei, were divided into three folds, each with six volumes for training and two for validation.

For the in-house data, the MaCS test set holds three volumes from the training acquisition sessions. The LSS set holds two crops of a spheroid that was never used for training. The ASD set holds 25 stacks of five aggregates, three HeLa Kyoto and two T-47D, each imaged at Z-steps of 0.5, 1, 2, 4 and 6 µm. In the benchmarks, ASD is represented by the stacks imaged at 2 µm, close to the Z-step of the training data, so that each specimen is counted only once; all Z-steps are used in the axial sampling analysis.

For the external datasets, the published test sets were kept unchanged (*Caenorhabditis elegans*, 7 volumes; BlastoSPIM 1.0 and 2.0 low-SNR, 60 and 3 volumes). NucMM releases only training and validation labels, so its published validation volumes served as the test set (NucMM-M, 4; NucMM-Z, 27). Fold 0 used each dataset’s own train/validation split. Folds 1 and 2 drew a validation set of the same size from the pooled training and validation volumes.

BlastoSPIM 1.0 is a time-lapse, with about eight frames per embryo taken 15 min apart. To keep these near-duplicate frames from inflating the training and validation sets, only one frame per embryo was kept (37 training and 18 validation embryos), and no embryo appears on both sides.

In total, each fold contained 150 training and 37 validation volumes. The decoder thresholds were tuned on the validation volumes only.

#### Isotropic resampling

For the general models, which combine images acquired with different imaging systems and voxel sizes, all volumes were resampled to isotropic voxel grids before training and inference. The target voxel size was selected for each dataset based on its typical nuclear diameter so that nuclei span approximately 24–26 voxels in every dataset, following an approach similar to that used in Cellpose. A voxel size of 0.5 µm was used for the in-house spheroid, BlastoSPIM and NucMM mouse datasets, where nuclei typically measure 11–14 µm in diameter. For the NucMM zebrafish and *C. elegans* datasets, which contain smaller nuclei with diameters of approximately 4 µm and 1.4 µm, isotropic voxel sizes of 0.17 µm and 0.06 µm were used, respectively. Images were resampled using trilinear interpolation and instance labels with nearest-neighbour interpolation. The in-house models were trained and applied on the native anisotropic grid, without resampling, and predictions of the general models on in-house data were mapped back to the native grid by nearest-neighbour interpolation for scoring.

### Compared methods

We compared our model with other published methods for 3D nucleus segmentation, each built on a different instance representation: star-convex polyhedra (StarDist-3D), diffusion flows (Cellpose-SAM), distance-field flows (Omnipose), and centroid-directed gradient fields (NucVerse3D). Each baseline method was trained with its authors’ published settings. For a fair comparison, every method ran for 70 epochs of 500 optimiser steps, at our batch size of 16, or the largest smaller batch that fits on a 48 GB GPU, and on our 24×192×192 patches wherever the method leaves the patch size free. Further method-specific settings are given and justified in the sections below.

### 3D U-Net

The 3D U-Net [15] is the volumetric extension of the well-established 2D U-Net [30]. An encoder progressively downsamples the input image into a low-resolution feature map, and a symmetric decoder reconstructs the full-resolution output from it. Skip connections concatenate the feature maps of the encoder with those of the decoder at each resolution level, so fine spatial detail lost during downsampling is preserved. The 3D U-Net was designed for semantic segmentation. Instance segmentation therefore requires an additional post-processing step, such as connected-component labelling or watershed.

### StarDist-3D

StarDist-3D represents each nucleus as a star-convex polyhedron [31]. For every voxel, the network predicts the probability of being the centre of a nucleus and the distances to the nucleus surface along a fixed set of radial rays, using a ResNet-based [32] or a U-Net-based CNN architecture. Instances are obtained by non-maximum suppression of the predicted polyhedra. Anisotropy is handled by distributing the rays according to the anisotropy of the objects and by subsampling the prediction grid only in XY. The representation requires star-convex objects, therefore, irregularly shaped nuclei cannot be described accurately.

StarDist-3D (StarDist 0.9.2, CSBDeep 0.8.2, TensorFlow 2.15.1) was trained from scratch with the settings of the 3D training example in the StarDist repository. The network was the default of StarDist’s 3D configuration, a U-Net backbone of depth 2 with 32 base filters. The publication used a ResNet [5] and reports very similar results with a U-Net. The Stardist-3D official repository uses U-Net as a backbone with 32 base filters. Volumes were normalised to their 1st–99.8th percentiles, holes in the labels were filled, and 96 rays were placed on a golden spiral adjusted to the object anisotropy of each fold’s training labels. In-house, the notebook’s grid of (1, 2, 2) failed its check that the network’s field of view covers the median training nucleus, so we used the smallest grid that passes, (1, 4, 4), as StarDist’s documentation advises for large objects; on the general split, the notebook’s rule, (2, 2, 2), passed. Training used the authors’ loss, optimiser, and augmentation at a batch size of 16 instead of 2, on our model’s 24×192×192 patches (general split 80×96×96), because 48×96×96 patches used in the original training notebook are thicker than our thinnest training volume. Validation used 64 fixed patches from the fold’s validation volumes instead of a random 15% of the training data; the learning rate was halved after 4,000 steps without improvement, the authors’ patience in steps, and the lowest-validation-loss weights were kept. The probability and NMS thresholds were optimised on the validation volumes with the authors’ routine over NMS values of 0.1–0.5.

### Cellpose

Cellpose [16] represents cells as flow fields. The network predicts the foreground probability and a flow field obtained by simulating heat diffusion from the centre of each cell, so the flows point toward the centre of the instance. Pixels are grouped into instances by following the flows until they converge at a common sink. The original model is based on a U-Net, while the recent Cellpose-SAM [33] replaces the encoder with a vision transformer. The network is two-dimensional: for volumes, Cellpose either averages the flows predicted on XY, XZ and YZ slices into a 3D flow field (3-axis mode), or segments each XY slice and stitches the masks across Z (2.5D mode).

Cellpose-SAM (cellpose 4.2.1.1, PyTorch 2.14.1) was fine-tuned on each fold from its released weights with the authors’ fine-tuning setup: AdamW with learning rate 10⁻⁵ and weight decay 0.1, batch size of 1, and the schedule of their 100-pass run (warm-up over the first tenth, halvings over the second half) compressed into the 35,000 steps. Training images were cut as the authors’ advice for 3D: XY, XZ, and YZ slices, with slices through Z stretched by the voxel aspect ratio, one random 512×512 crop per slice, and crops with fewer than five nuclei discarded. We took 100 slices per axis and volume in-house instead of the authors’ 10, because our six completely annotated training volumes would otherwise give only a few crops, but 10 on the general split. Crops were normalised to their 1st-99th percentiles and trained with the library’s loss and augmentation on 256×256 tiles; the final weights were used, as in the authors’ training function. In the 3-axis mode, each volume’s voxel aspect ratio was passed to Cellpose’s own anisotropy parameter. In the 2.5D mode, each XY slice was segmented in 2D, and masks on consecutive slices were joined into one 3D object when their intersection over union was at least 0.25, the default of Cellpose’s stitching function. 3D flow smoothing was left at the library default.

### Omnipose

Omnipose [17] builds on Cellpose but derives the flows from the gradient of a distance field instead of a diffusion from the cell centre. The flows therefore converge on the medial axis (skeleton) of each object instead of a single centre point. This makes the method better suited to elongated, branched and irregularly shaped cells, for which a single centre is poorly defined. For near-spherical cells, the skeleton approaches a single point, and the flows become similar to those of Cellpose. Unlike Cellpose, Omnipose can be trained as a native 3D network, which has no correction for anisotropic voxels.

Omnipose (version 1.1.4, PyTorch 2.14.1) was trained from scratch, following the authors’ published 3D training command. We used the library’s default optimiser, SGD (learning rate 0.1, momentum 0.9, weight decay 10⁻⁵), together with its built-in schedule: a short linear warm-up, a constant rate, and a stepwise decay over the final 2.5% of training. Because this schedule is defined for the authors’ 4,000 epochs, we stretched it proportionally over our 35,000 training steps. The network predicted the flows and the distance field. Crops were 96×96×96 voxels, the voxel count of our patches, instead of the default 128³, at batch 2, because the command’s batch of 4 did not fit in memory. Because Omnipose rescales the whole image before every crop, training volumes were normalised to their 1st–99th percentiles and cut into blocks of at most 256³ voxels. Training used the authors’ augmentation, and the final weights were used for testing. The mask threshold was selected during validation on the validation volumes from the range −3 to 2.

### NucVerse3D

NucVerse3D [20] is a recent method for nucleus segmentation across microscopy modalities. It uses a 3D residual attention U-Net to predict the foreground and a centroid-directed gradient field along the X, Y, and Z axes, similarly to HoVer-Net in 2D [34]. To reconstruct the instances, each foreground voxel is moved iteratively against the predicted gradient field. The converged positions are clustered into nucleus centroids, and each voxel receives the label of the centroid it reached.

NucVerse3D (commit d809a2e; TensorFlow 2.16.2, Keras 3.12.4) was trained from scratch with the authors’ code and network as their unscaled generalist variant, which works on the native voxel grid without anisotropy correction. Volumes went through the authors’ preprocessing at training and inference. Patches of 64×128×128 voxels were drawn with the authors’ patch builder, 800 for training and 200 for validation per dataset. On the general split, each source dataset was an individual dataset, as the authors do, whereas in-house, each volume was, which weights every volume equally regardless of its size. Validation patches came from the fold’s validation volumes, while training patches were stored in random order across volumes. Training used the authors’ augmentation, loss and optimiser (Adam, learning rate 10⁻³) at batch 4 instead of 2, kept the authors’ learning-rate halving and early stopping at their patience of 10 and 15 passes, and kept the lowest-validation-loss weights. Instances were reconstructed with the authors’ tiled prediction and gradient-descent clustering at their default settings.

### CellTouch

#### Network architecture

CellTouch has a 3D U-Net [15] as a backbone. The encoder has six resolution levels with 32, 64, 128, 256, 320 and 320 feature maps. Each level contains two convolutions, each followed by instance normalisation and a leaky ReLU activation. A decoder restores the full resolution with transposed convolutions, and skip connections concatenate the encoder feature maps into the decoder at each level. Because the volumes of our in-house datasets are sampled 3.5 to 26 times more coarsely along the Z-axis than in the XY, we use an anisotropic pooling schedule. The first two levels pool only in the XY plane (1×2×2), and the remaining three levels pool in all directions (2×2×2), for a total stride of 8×32×32 (Fig. 2). Following nnU-Net [35], convolutions are 1×3×3 in the first three levels, while the Z-axis has not yet been pooled, and 3×3×3 after. This preserves the axial resolution in the shallow layers, where a nucleus spans only a few planes [35].

**Fig. 2.**
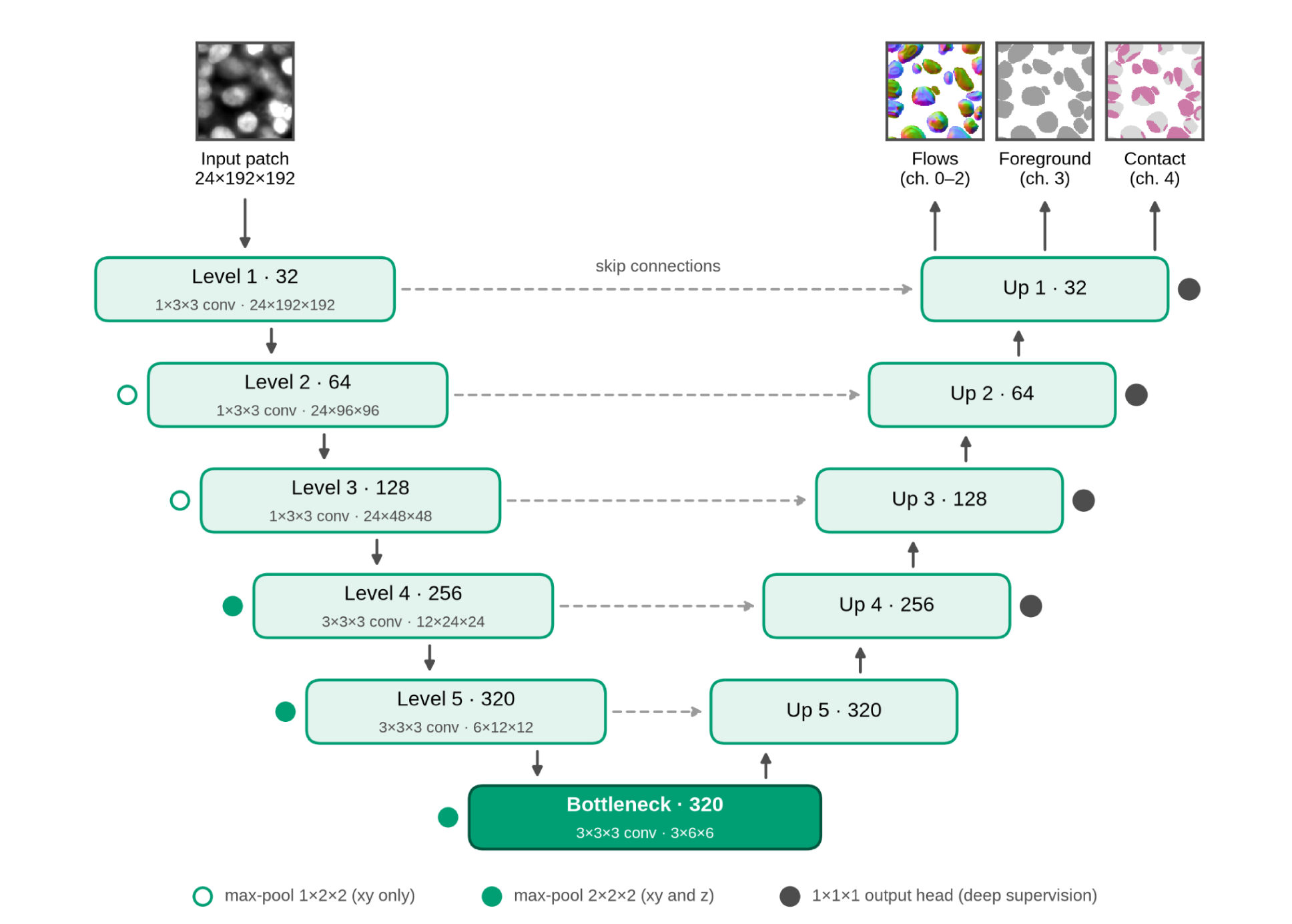
Architecture of the CellTouch Network. The backbone is a 3D U-Net that receives an input patch and predicts five output channels: channels 0–2 are the flow field, one component per axis, channel 3 is the nuclear foreground, and channel 4 is the contact channel.

#### Outputs

CellTouch predicts five output channels from the final decoder layer (Fig. 2). Three channels form a flow field, one component per axis, defined as the normalised gradient of the distance to the nucleus surface and pointing toward the interior of each nucleus, as in Omnipose. One channel is the nuclear foreground, and the last is the contact channel, which marks where two nuclei touch. Instances are built from the foreground and the contact channel alone: the contact surface is removed from the foreground, so touching nuclei fall apart into separate components.

#### Contact channel

The contact channel is a binary target that marks only the surfaces where two nuclei touch and excludes surfaces facing the background. Related signals have been used in 2D images as a loss weight on the gaps between touching cells [30] or as a separate touching-border class [36], whereas here the contact surface is defined in 3D and scaled to the voxel anisotropy. It is derived from the instance labels in two steps. First, contact voxels are identified as foreground voxels whose 3 × 3 × 3 neighbourhood contains at least two different nucleus labels. This is computed with a maximum and a minimum filter over the label volume, in which background voxels are excluded from the minimum, so that a nucleus bordering only background is not counted as a contact. The neighbourhood spans three planes in z, so nuclei touching along the axial direction are detected as well as nuclei touching side by side. Second, the contact surfaces are dilated into a shell: a Euclidean distance transform is computed from the contact voxels, with the axial distance weighted by the volume’s axial-to-lateral sampling ratio. Every foreground voxel within 3.5 lateral pixels of a contact is labelled as contact. Weighting by the sampling ratio makes the shell equally thick in physical units along z and in-plane within a volume, so a coarse Z-step doesn’t make it artificially thick across planes. Voxels outside nuclei are never labelled as contact. The resulting shell covers about 10% of nuclear voxels in our volumes. The network is trained to predict it with a binary cross-entropy loss weighted by ×20 for the positive class. At inference, the predicted shell is removed from the predicted foreground, the remaining connected components are labelled as nuclei, and the removed voxels are reassigned to the nearest component (carving).

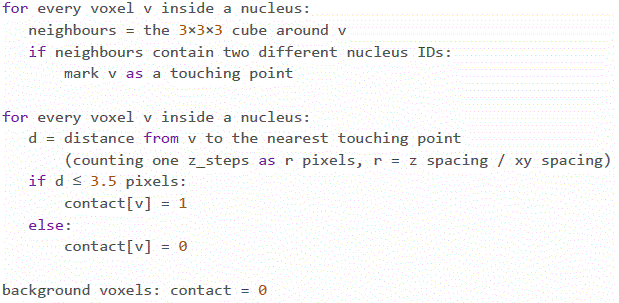

#### Loss function

CellTouch is trained with three loss terms: binary cross-entropy on the foreground channel, binary cross-entropy on the contact channel, and mean squared error on the flow field. The contact term uses a positive-class weight of 20, because contact voxels make up only about 1% of a training patch and an unweighted loss is minimised by predicting no contact anywhere. Following Cellpose, the flow targets are multiplied by 5 so that the regression term is on the same scale as the classification terms. The flow error is computed only on foreground voxels, since background flows are zero by construction, and including them would reward an empty prediction. All three terms have weight 1. The network is trained with deep supervision: decoder level *l* (*l* = 0 at full resolution) receives weight α_l = 0.5^l, the coarsest level is excluded, and targets are downsampled to each level by nearest-neighbour sampling. The total loss is the weighted mean of the foreground, contact and flow losses.

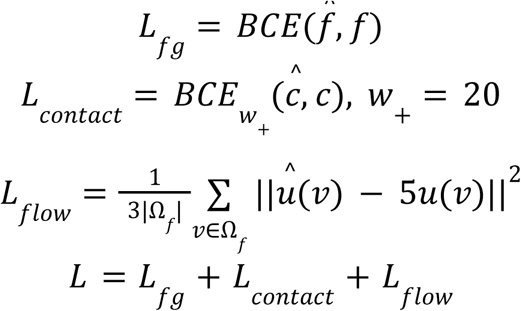

#### CellTouch training

Each model was trained in each of three folds, using six training and two validation volumes per fold. Models were trained on patches of 24×192×192 voxels with a batch size of 16 for 70 epochs of 500 iterations, and half of the patches were sampled to contain foreground. We used stochastic gradient descent with Nesterov momentum of 0.99, an initial learning rate of 3×10⁻³ with polynomial decay, and weight decay of 3×10⁻⁵. Training data were augmented with random flips and in-plane 90° rotations, with the flow vectors transformed consistently. Intensities were clipped at the 0.5th and 99.5th percentiles and normalised to zero mean and unit variance for each volume. Training used mixed precision on a single NVIDIA RTX A6000 GPU and took about 7 hours per model. The model after the final epoch was used for evaluation; validation was used only to monitor training.

#### CellTouch inference

Volumes were predicted with a sliding window of the training patch size and 25% overlap between windows, using the same intensity normalisation as in training. For the axial-resolution series, where nuclei occupy only about 0.25% of the volume, intensities were clipped at the 99.99th percentile to avoid saturating the nuclei. The predicted channels were decoded with the native decoder of each configuration, using the anisotropy of each volume for distance computations. Test-time augmentation was not used for the reported results. Each fold was evaluated separately, and results are reported as the mean over the three folds.

### Evaluation metrics

#### F1 score

Predicted nuclei were matched one-to-one to ground-truth nuclei by their intersection over union (IoU; Eq. 1). At an IoU threshold t, a predicted nucleus was counted as a true positive (TP) when it was matched to a ground-truth nucleus with IoU > t. Predicted nuclei without such a match were counted as false positives (FP), and ground-truth nuclei without one were counted as false negatives (FN). Above t = 0.5 every match is unique; at t = 0.5 the matching was chosen to maximise the total IoU. The F1 score was computed from these counts at each threshold (Eq. 2). F1 is reported at t = 0.5, which mainly measures detection, and at t = 0.75, which also requires accurate outlines. The mean F1 over nine thresholds from 0.5 to 0.9 in steps of 0.05 is also reported (Eq. 3). Scores were computed per test volume, without test-time augmentation. They were averaged over the volumes of a test set and over the three cross-validation folds and then over test sets, with each set weighted equally.

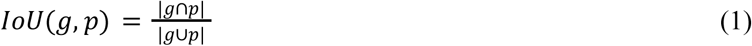

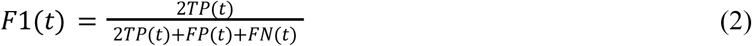

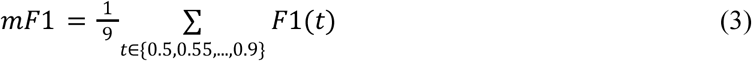

#### Panoptic quality

Panoptic quality (PQ) [37], [38], [39] combines detection and delineation in a single number (Eq. 4). It is the product of the recognition quality (RQ) (Eq. 5) and the segmentation quality (SQ) (Eq. 6). RQ is computed from the numbers of true positives (TP), false positives (FP), and false negatives (FN). SQ is the mean intersection over union (Eq. 1) of the matched pairs of ground-truth and predicted nuclei. A pair was matched when its IoU exceeded 0.5, a threshold at which every match is unique. RQ therefore measures how many nuclei are found correctly, and SQ how accurately the found nuclei are delineated.

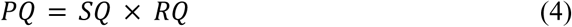

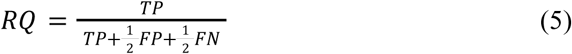

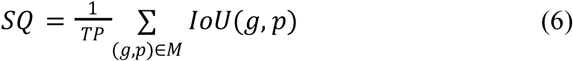

#### Aggregation

All metrics were computed per volume and then averaged over the volumes of each test set; objects were not pooled across volumes. Each method was evaluated separately in each of the three folds, and we report the mean over folds. In addition, we report the number of ground-truth nuclei found (matched at IoU 0.5) and the total number of predicted objects, which shows over– and under-segmentation directly. During training, the mAP (Eq. 7, 8) was computed every 10 epochs on the validation volumes to monitor convergence. To reduce memory use and evaluation time, it was computed on a 48×512×512 crop from the densest region of each validation volume. The validation mAP was used only for monitoring, not for model selection: the model after the final epoch was used for all reported results.

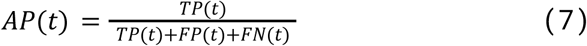

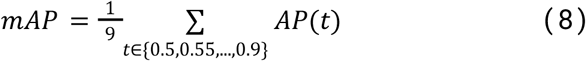

## Results

As 3D nuclei segmentation was previously introduced as a crucial part of the image-based 3D-oid analysis platform, here we aimed to improve the segmentation performance to allow a better prediction tool for diagnostic purposes [6]. Therefore, the training dataset included a high variety of publicly available 3D-oid models screened with light-sheet fluorescence microscopy, providing high-quality single-nucleus resolution in dense samples. To provide a fair comparison of the segmentation performance, unseen datasets from diverse 3D specimens were selected for testing.

CellTouch adds to the network output a contact channel that marks only the surfaces where two nuclei touch (Fig. 2). We first tested whether this channel drives the improvement by training the same network with and without it on four different target representations before comparing CellTouch with state-of-the-art methods and measuring its sensitivity to axial sampling.

### Ablation study

To test the effect of the contact channel, we combined it with the output representations of four popular segmentation methods. The backbone network for the study was the 3D U-Net (Fig. 2). The first representation is the binary foreground. The second one is a three-class map that predicts the background, the nucleus interior, and its boundary surface. The third one predicts, for every voxel of a nucleus, the Euclidean distance to the nucleus surface. The last is a three-dimensional flow field, defined as the normalised gradient of the distance map and pointing toward the interior of each nucleus, as in Omnipose. Each representation was trained with and without the contact channel, a thin shell of constant physical thickness that marks only the surfaces where two nuclei touch and excludes their outer surfaces. The contact channel improved every representation on all measured metrics (Fig. 3). It mainly improved detection quality, increasing F1 at IoU 0.5 by 0.15 to 0.38. The improvement in F1 at IoU 0.75 was smaller, 0.04 to 0.15, which implies it helps with better object recognition rather than segmentation.

**Fig. 3:**
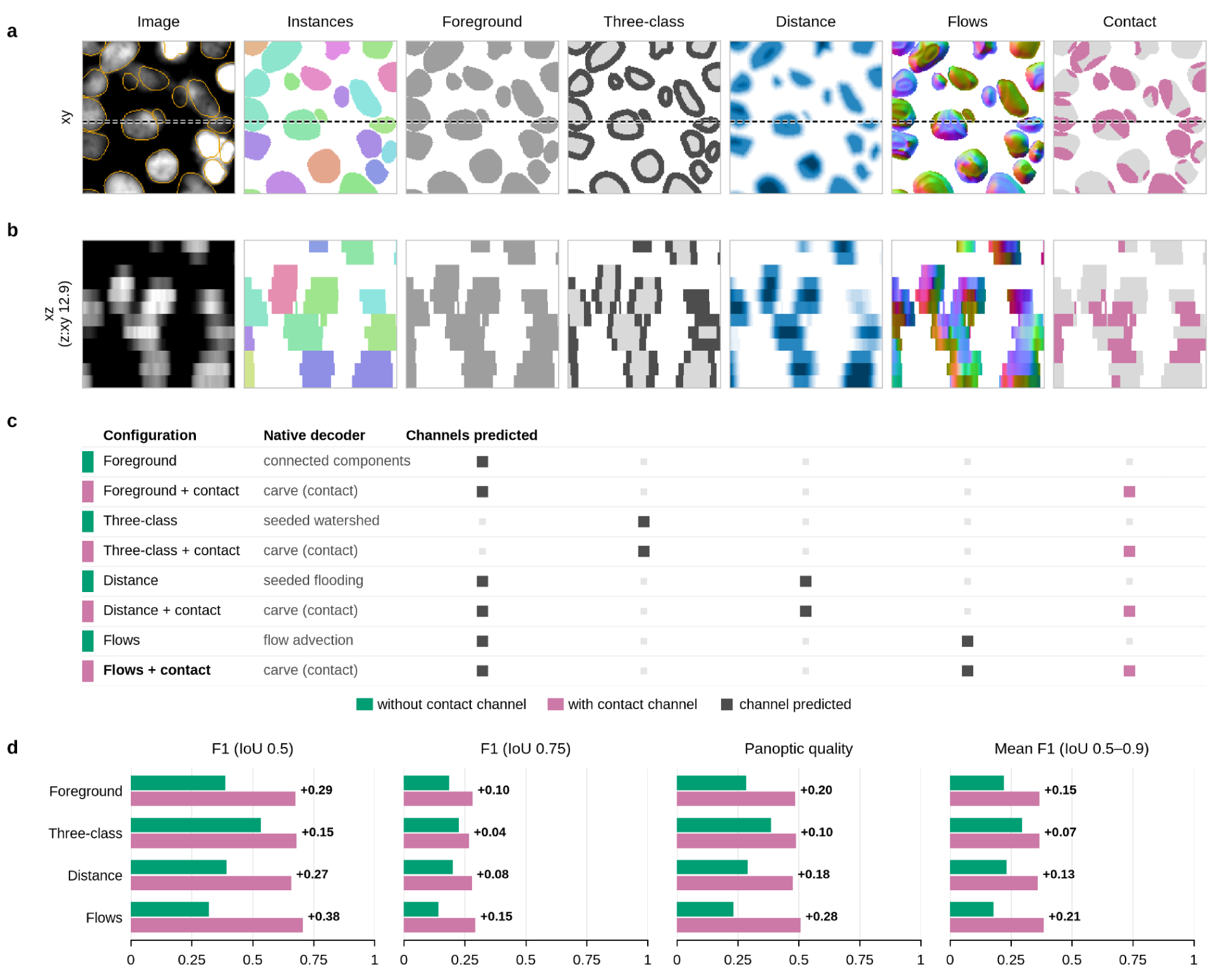
Contact-channel ablation across target representations. **a, b**. Representative example in a lateral xy plane (a) and an xz section along the dashed line (b), showing the image and each representation: instances, each nucleus in a different colour; foreground; three classes, background, foreground and boundary; distance to the nucleus boundary, highest at the centre; the three-component flow field; and the proposed contact channel. (c), The eight combinations, the channels each one predicts, and its native decoder. (d), F1 at IoU 0.5 and 0.75, panoptic quality and mean F1 over IoU 0.5–0.9 on the MaCS test set for each representation trained without (green) and with (magenta) the contact channel; the number next to each pair is the gain. Values are averaged over the three cross-validation folds.

### Benchmarks

We benchmarked CellTouch against four state-of-the-art methods for 3D nucleus segmentation: StarDist-3D, Cellpose-SAM with 2.5D and with 3D flow decoding, Omnipose, and NucVerse3D. Each method was trained in two settings: on our anisotropic in-house data alone and on a general collection that pools the in-house data with five public datasets. In both settings, every method was trained or fine-tuned on the same training volumes with its own validation-based settings, and accuracy was measured as the mean F1 score over IoU thresholds from 0.5 to 0.9, alongside F1 at IoU 0.5 and 0.75 and panoptic quality. On the in-house data, the methods were tested on three held-out sets: Mono and Co-culture Spheroids, Large Spheroid Subregions, and the Axial Sampling Dataset imaged at a 2 µm z-step (Fig. 4). CellTouch reached the highest mean F1 on all three test sets (Fig. 4b) and the highest mean over the sets, 0.557. StarDist-3D followed with 0.422, then Cellpose-SAM 2.5D with 0.359, Cellpose-SAM 3D with 0.273, and NucVerse3D with 0.120 (Fig. 4a). Omnipose produced almost no correct segmentations on the anisotropic volumes, with a mean F1 of 0.002. The difference was largest at strict overlap: at IoU 0.75, CellTouch reached an F1 of 0.559 against 0.339 for StarDist-3D, whereas at IoU 0.5 the two were close at 0.799 and 0.778. CellTouch therefore detected about as many nuclei as StarDist-3D but delineated them more accurately, which also gave it the highest PQ, 0.630 against 0.560. Compared volume by volume, CellTouch had a higher mean F1 than every competitor on all ten test volumes (two-sided Wilcoxon signed-rank test, p = 0.002, the smallest value attainable with ten volumes, Holm-corrected p = 0.01).

**Fig. 4:**
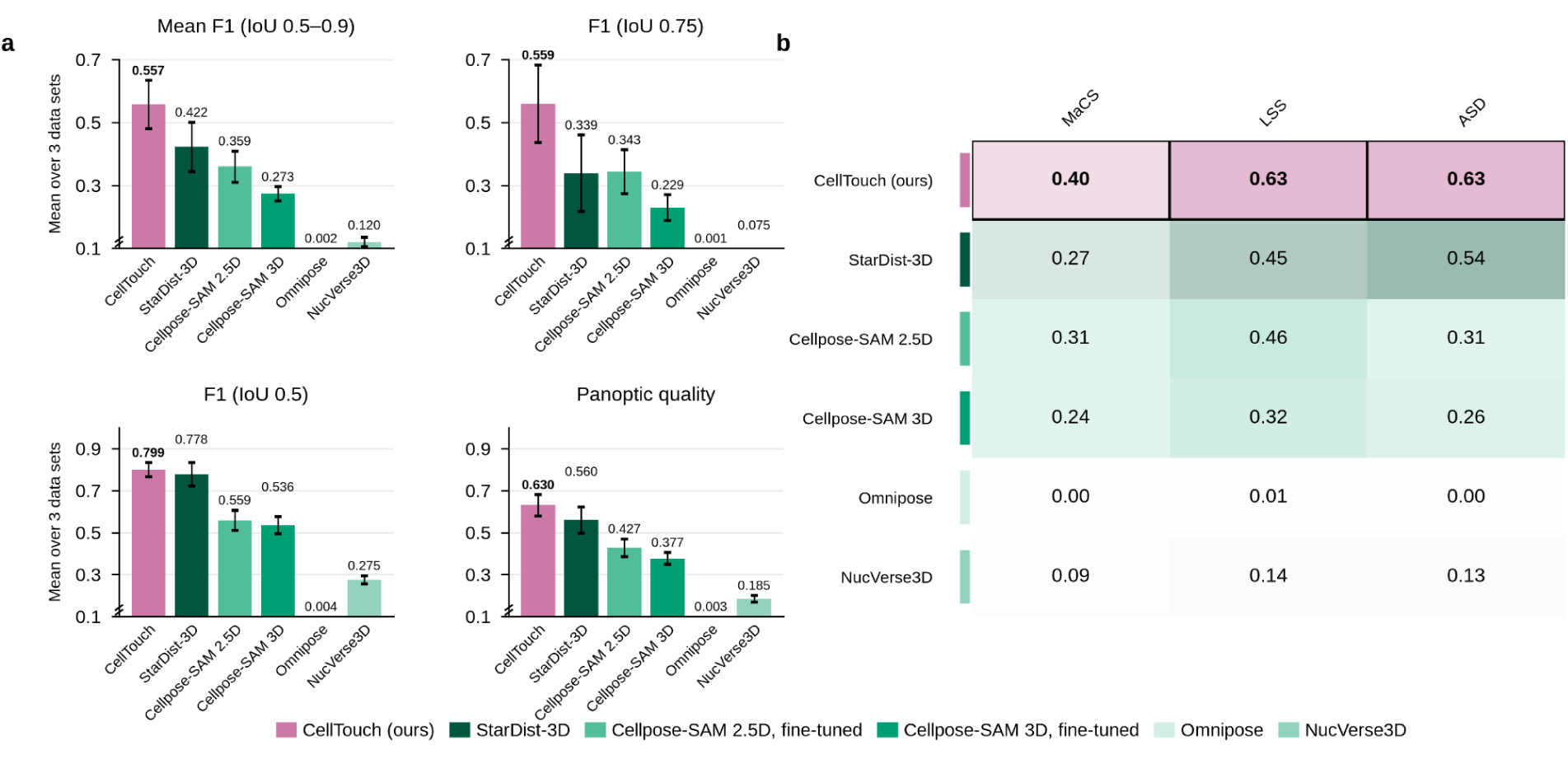
Benchmark models trained on the anisotropic in-house data. (a) Mean F1, F1 at IoU 0.75 and 0.5, and panoptic quality, averaged over three folds and over the three in-house test sets. (b) Mean F1 per test set, with the best method per set outlined.

### External dataset benchmark

In the general setting, the in-house training volumes were pooled with the training splits of BlastoSPIM 1 and 2, NucMM-Z, NucMM-M and *C. elegans*. The methods were then tested on all eight test sets, 111 volumes in total (Fig. 5). CellTouch reached the highest mean F1 over the eight sets, 0.580. Cellpose-SAM 3D followed with 0.515, then StarDist-3D with 0.508, NucVerse3D with 0.500, Cellpose-SAM 2.5D with 0.397 and Omnipose with 0.274 (Fig. 5a). CellTouch performed best on seven of the eight sets (Fig. 5b). The exception was NucMM-M, where nuclei rarely touch and StarDist-3D reached 0.63 against 0.57. As on the in-house data, the advantage came from more accurate outlines. At IoU 0.75, CellTouch reached an F1 of 0.598, against 0.494 for Cellpose-SAM 3D, the best competitor, and 0.442 for StarDist-3D. At IoU 0.5, StarDist-3D was marginally ahead, at 0.873 against 0.865 (p = 0.068). Compared volume by volume, CellTouch had a higher mean F1 than StarDist-3D on 93 of 111 volumes, than Cellpose-SAM 3D on 94 and than NucVerse3D on 100, all with p < 10⁻¹¹. It was higher than Cellpose-SAM 2.5D on 108 volumes and than Omnipose on all 111 (p < 10⁻¹⁸; two-sided Wilcoxon signed-rank test, Holm-corrected over the five competitors). CellTouch also had the highest PQ, 0.672 against 0.643 for StarDist-3D (p < 10⁻⁷). Our model’s advantage was not specific to our own data, on the five public data sets alone, CellTouch reached a mean F1 of 0.594 against 0.554 for StarDist-3D, the best competitor there.

**Fig. 5:**
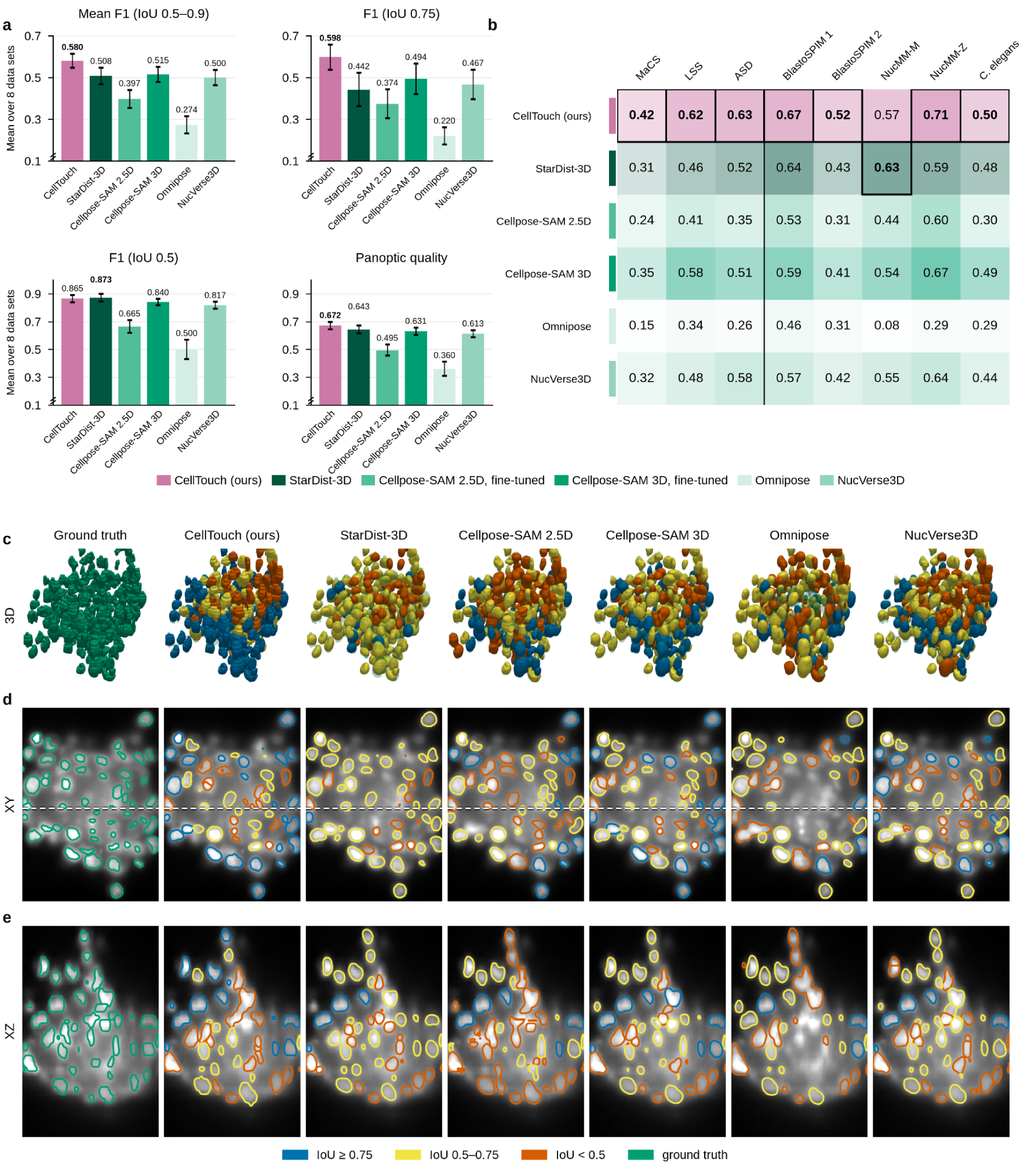
**a-b** Benchmark on pooled external datasets (a) Mean F1, F1 at IoU 0.75 and 0.5, and panoptic quality, averaged over three folds and over eight datasets (111 volumes); CellTouch outperforms every competitor in mean F1, F1 at IoU 0.75, and panoptic quality (per-volume Wilcoxon, Holm p < 10⁻⁶). (b) Mean F1 per dataset; the vertical line separates in-house from public sets. **c-e**: A co-culture spheroid in 3D (c), with XY (d) and XZ (e) sections through the same window, segmented by the general-trained models. Each predicted nucleus is coloured by its best IoU with the ground truth, shown as glass in (c) and coloured outlines in (d) and (e), and surfaces are smoothed along z for display. CellTouch delineates the most nuclei at IoU ≥ 0.75.

The difference in outline accuracy is visible in the segmentations (Fig. 5c–e). StarDist-3D placed nearly every nucleus correctly, but most of its matches fell between IoU 0.5 and 0.75, and only 17 of the 265 nuclei exceeded 0.75. Cellpose-SAM in 2.5D mode fragmented nuclei along the Z-axis, as each plane was segmented separately and stitched afterwards, while Omnipose merged neighbouring nuclei into elongated objects. CellTouch matched 120 nuclei at IoU ≥ 0.75, against 73 for Cellpose-SAM 3D, the next best method. Its remaining errors, like those of every method, were concentrated in the core of the spheroid, where the nuclear signal was several-fold weaker than at the periphery.

### Axial sampling robustness analysis

Based on our anisotropic data, we compared CellTouch to StarDist-3D to analyse its sensitivity to anisotropic data. While on F1@0.5 the scores are nearly identical, 0.94 for both CellTouch and StarDist-3D at a Z-step of 2 µm, on F1@0.75 the gap between CellTouch and StarDist-3D increases as the data become less isotropic, from 0.12 at 0.5 µm to 0.29 at 2 µm and 0.28 at 4 µm (Fig. 6a). At 6 µm, where a nucleus spans only two to three sections, both methods fall below 0.06 at this threshold. Models trained on anisotropic grids are more sensitive to different Z-steps (Fig. 6b): StarDist-3D trained on our in-house data failed almost completely at 0.5 µm, with an F1@0.5 of 0.03, while CellTouch trained on the same data kept an F1@0.5 of 0.75. Both in-house models performed best close to the anisotropy of their training data and lost accuracy at coarser Z-steps.

**Fig. 6.**
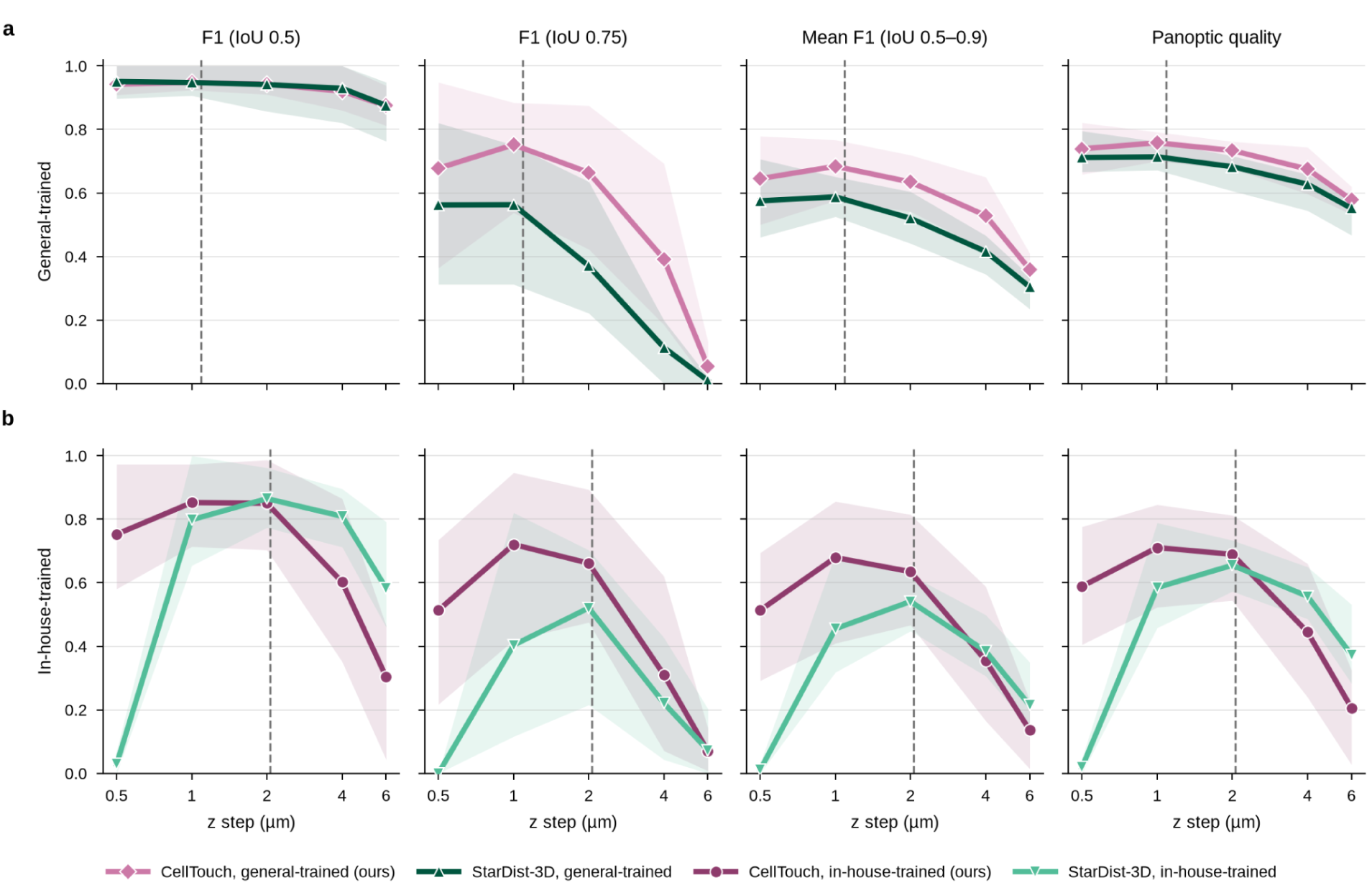
Sensitivity to axial sampling. F1 at IoU 0.5 and 0.75, mean F1 over IoU 0.5–0.9 and panoptic quality against Z-step for CellTouch and StarDist-3D trained on **a**, the general data, and **b**, our in-house data. Lines show the mean of five specimens, three HeLa Kyoto and two T-47D aggregates, each imaged at all five Z-steps. Dashed lines mark the mean anisotropy of the training volumes.

## Discussion

As modern microscopy platforms develop to acquire highly detailed volumetric datasets from 3D-oids in a high-throughput manner, 3D data has become essential for investigating such complex systems. These technological developments provide opportunities to investigate cellular organisation, morphology, and interactions within physiologically relevant environments. However, the increasing volume and complexity of image data place substantial demands on downstream analysis workflows.

Many existing analysis pipelines still rely on computationally efficient 2D segmentation strategies applied to inherently 3D structures [13], [16], [40]. However, such slice-by-slice analysis of 3D datasets often fails to accurately represent object morphology across the imaging volume, particularly when objects overlap or extend across multiple z-planes. To address these challenges, robust 3D segmentation tools capable of operating directly on volumetric data are continuously developed to perform consistently under a wide range of imaging conditions, including variations in signal intensity, image quality, noise levels, and anisotropic z-resolution.

In this context, CellTouch represents a significant methodological advancement by exploiting the contact representation of touching objects to separate them. In the case of 3D-oid models, separating densely arranged, occasionally touching nuclei represents a major challenge, and it becomes harder as the Z-axis is sampled more coarsely. By learning where neighbouring nuclei touch, CellTouch separates them directly from the network output.

Experimenting with the contact channel was motivated by a simple observation, while we noticed that the boundary predictions were not strong enough to separate two or more touching nuclei. The boundary class was predicted confidently at the outer surfaces but only weakly where nuclei touched. One of the main reasons is class imbalance. Outer surfaces make up about 98% of all nucleus surfaces and coincide with a drop in intensity, so a network learns them readily, whereas the internal interfaces between touching nuclei are rare and often show little contrast, yet they decide whether a touching pair is split or merged.

We conducted an ablation study to test whether the contact channel truly helps and found that it naturally improves nucleus detection compared with the baseline representations, as it directly targets the interfaces between touching nuclei that decide whether they are split or merged.

Across eight data sets spanning spheroids, aggregates, embryos, *C. elegans* and electron microscopy, it delineated nuclei significantly more accurately than StarDist-3D, Cellpose-SAM, Omnipose and NucVerse3D, with a mean F1 of 0.58 against 0.52 for the strongest competitor, Cellpose-SAM 3D (per-volume Wilcoxon, p < 10⁻¹⁴). The advantage lay in outline accuracy, not in detection: F1 at IoU 0.75 was 0.60 against 0.49 for the strongest competitor at this threshold, Cellpose-SAM 3D, while detection at IoU 0.5 was on par. Removing the contact channel showed that this channel accounts for the gain, raising the share of merged touching nuclei from 8% to 84%. In densely packed, anisotropically sampled cultures, where current methods merge neighbours or fail, CellTouch was the most accurate method.

The same design made CellTouch robust to the Z-step size. On five specimens of two cell lines, each imaged at Z-steps from 0.5 to 6 µm, CellTouch matched StarDist-3D in detection, and its advantage in outline accuracy grew as the data became less isotropic. Models trained on the native anisotropic grid of our in-house data were considerably more sensitive to the Z-step size: CellTouch trained on the same data remained usable across a wider range than StarDist-3D, which lost almost all nuclei at 0.5 µm. At 6 µm, where a nucleus spans only two to three sections, outline accuracy dropped for both methods. Sampling at 2 µm instead of 0.5 µm costs little accuracy while acquiring four times fewer sections. Coarse z sampling therefore remains compatible with accurate single-nucleus segmentation, saving acquisition time, light exposure and storage in 3D screening. Extending the anisotropic benchmark to further cell lines and microscopes and confirming that downstream nuclear features are preserved across Z-steps are the next steps towards routine use.

Such precise detection integrated with automated analysis pipelines may enable large-scale, reproducible quantification of complex biological systems and will facilitate advances in phenotypic screening, developmental biology, and tissue engineering. As imaging technologies continue to evolve, the development of robust, generalisable 3D segmentation approaches will be essential for fully exploiting the information contained within next-generation volumetric datasets. Looking ahead, the impact of CellTouch is expected to extend beyond microscopy image analysis, as accurate volumetric object identification and separation showed promising results.

## Data availability

The in-house MaCS, LSS and ASD datasets, with annotations, are available at https://doi.org/10.5281/zenodo.24563346. Nuclei were annotated in napari with the Napari-nD-annotator plugin (Hollandi et al. [24]; https://github.com/bauerdavid/napari-nD-annotator) by drawing contours in 2D slice by slice. The MaCS volumes include previously published co-culture spheroids (https://doi.org/10.6084/m9.figshare.24982227) and optically cleared spheroids (https://doi.org/10.6084/m9.figshare.12620078). The public datasets are available from their original sources: BlastoSPIM 1.0 and 2.0 (https://doi.org/10.25452/figshare.plus.26540593), NucMM (https://nucmm.grand-challenge.org/) and *C. elegans* (https://doi.org/10.5281/zenodo.5942575).

## Code availability

The maintained CellTouch package is available at https://github.com/grexai/CellTouch. The code to reproduce all analyses and figures of this study is available at https://github.com/grexai/CellTouch-paper. Trained CellTouch weights are available at https://huggingface.co/grexai/Celltouch, and the weights of the competing methods retrained on our data splits at https://huggingface.co/grexai/celltouch-competitors.

## Acknowledgements

I.G., A.D., M.B., C.M., E.R., and P.H. acknowledge generous support from the Lendület BIOMAG grant (no. 2018–342), TKP2021-EGA09 (TKP-31-8/PALY-2021), Horizon-BIALYMPH (2023-1.2.1-ERA_NET-2023-00002), Horizon-SYMMETRY (2019-2.1.7-ERA-NET-2022-00044), Horizon-SWEEPICS (101135053), H2020-Fair-CHARM (101016457), HUNRENTECH (TECH-2024-34), HAS-NAP3 (NAP2022-I-6/2022), TKCS-2024/73, 2026-4.1.1-MISSZIÓK-2026-00025; PerMel-AI (EPPERMED2024−590) and grants from OTKA-SNN no. 139455/ARRS and NKKP (ADVANCED_24_151202). P.H. gratefully acknowledges support from the FIMM High Content Imaging and Analysis Unit (FIMM-HCA; HiLIFE-HELMI) and the Finnish Cancer Society.

## Contributions

Conceptualisation: I.G., A.D., P.H.;

Data curation: E. R., A. D.;

Formal analysis: I.G., E. R., A. D.;

Funding acquisition: A.D., P.H.;

Investigation: I.G.;

Methodology: I.G., A.D.;

Project administration: A.D.;

Resources: A.D., P.H.;

Software: I.G., M.B.;

Supervision: I.G.;

Validation: I.G., M.B.;

Visualisation: I.G.;

Writing – original draft: I.G., E. R., M.B.;

Writing – review & editing: I.G., A.D., P.H.;

